# Pro-cognitive reshaping of neuronal dynamics by a human CSF-based factor

**DOI:** 10.1101/2024.12.26.629959

**Authors:** Marc Dos Santos, Marc P. Forrest, Ewa Bomba-Warczak, Euan Parnell, Seby L. Edassery, Kun Yang, Lindsay N. Hayes, Jennifer M. Coughlin, Blair L. Eckman, Catherine Lammert, M. Dolores Martin-de-Saavedra, Maria Barbolina, Akira Sawa, Jeffrey N. Savas, Peter Penzes

## Abstract

Neuronal connection dysfunction is a convergent cause of cognitive deficits in mental disorders. Cognitive processes are finely regulated at the synaptic level by membrane proteins, some of which are shed and detectable in patients’ cerebrospinal fluid (CSF). However, whether these soluble synaptic proteins can harnessed as innovative pro-cognitive factors to treat brain disorders remains unclear. Here, we use quantitative proteomics to identify shed synaptic proteins dysregulated in the CSF of subjects with schizophrenia (SCZ), a mental disorder characterized by cognitive and synaptic dysfunction. The level of a yet uncharacterized soluble form of the voltage-gated calcium channel auxiliary subunit, α2δ-1, is robustly reduced in SCZ CSF. Remarkably, soluble α2δ-1 is convergently downregulated across several brain disorder CSF proteomes. We show that the brain releases soluble α2δ-1 in an activity-dependent manner, which can reorganize neuronal network dynamics by binding to synaptic targets and promoting inhibitory neuron plasticity. A single brain injection of a synthetic soluble α2δ-1 improved interneuron and cognitive deficits in a mutant mouse model of SCZ and cortical dysfunction. These findings underscore the potential of shed synaptic proteins as novel therapeutic agents capable of enhancing brain function in diverse brain disorders characterized by cognitive impairment.

## Main Text

Cognitive deficits are a series of symptoms shared by numerous brain disorders ranging from neurodegenerative^1^ to neuropsychiatric diseases ^2^ and these symptoms are among the most challenging to treat. For example, the antipsychotic drugs used to treat schizophrenia (SCZ), a severe chronic psychiatric disorder, usually fail to address its cognitive symptoms^3–5^. While multiple factors contribute to SCZ, synaptic abnormalities in excitatory and inhibitory neurons have emerged as critical players in the pathophysiology of many psychiatric disorders^6–8^. Large-scale human genomic studies have revealed that SCZ is highly heritable and that many risk genes for psychiatric disorders encode synaptic proteins, many of which are membrane-bound^9,10^. Some membrane proteins undergo ectodomain shedding, in which proteases, called sheddases, cleave the protein to release a soluble extracellular protein fragment that can have transcellular signalling properties^11–14^. Many of these ectodomains are detectable in the cerebrospinal fluid (CSF), representing the neuronal sheddome, which is actively interfaced with the brain’s extracellular milieu^11,14^. Recently, several studies have reported that proteins detected in the CSF^15^ and blood^16,17^ have pro-cognitive properties in aged mice, supporting the possibility that CSF-derived soluble factors could have therapeutic benefits for mental disorders. Therefore, we asked whether the synaptic sheddome is altered in SCZ and if some of the candidate proteins were convergently dysregulated in neuropsychiatric disorders linked to synaptic and cognitive deficits, such as Alzheimer’s disease (AD), aging, bipolar disorder (BIP) and major depressive disorders (MDD)^18–20^. CSF proteome analysis revealed that CACNA2D1, encoding the auxiliary subunit α2δ-1, previously unknown to undergo shedding, was reduced in SCZ CSF. Cross-disorder proteome re-analysis of several published CSF studies of brain disorders showed that α2δ-1 was also reduced in other brain disorders affecting cognition. We observed that neuronal activity and environmental enrichment, two processes linked to cognitive performance, increased soluble α2δ-1 released in mice. We designed a recombinant form of soluble α2δ-1 (SEAD1) and found that it bound preferentially to PV+ interneurons and their synapses, interacting with both excitatory pre- and postsynaptic proteins. Functionally, SEAD1 modulated neuronal network dynamics in a GABAergic neuron-dependent manner. Finally, injecting SEAD1 normalized the interneuron and behavioural phenotype of a mouse model of 16p11.2 duplication syndrome, a neurodevelopmental condition associated with SCZ and cortical network alterations^21,22^. Together, these results support the potential of synaptic ectodomains detected in human CSF to yield new diagnostic tools and therapies for neuropsychiatric and cognitive disorders.

### Reduced α2δ-1 protein levels in the CSF in schizophrenia and other cognitive disorders

To explore how the synaptic sheddome is affected in SCZ, we used multiplexed tandem mass tag (TMT) mass spectrometry-based quantitative proteomics^23^ to measure the relative abundance of proteins in CSF samples from individuals with SCZ and unaffected controls (UC) (**Fig. 1a**). Our analysis identified over 1400 unique proteins and revealed reduced levels of several synaptic proteins in SCZ CSF samples compared to UC (**Fig. 1b and Supplementary Information Table 1**). First, using the Cell-type Specific Expression Analysis (CSEA) tool, we observed that although total CSF proteins are broadly expressed in the brain (**Extended data Fig 1a**), differentially expressed proteins (DEPs) in SCZ are enriched in genes putatively originating from the neocortex, a brain region strongly implicated in cognitive processes and SCZ (**Fig. 1c**)^7,24-27^. We prioritized important proteins for brain function based on high evolutionary constraint (pLI > 0.9) from gnomAD^28^ (Table S2). Notably, the CSF proteome showed modest enrichment of these genes (enrichment ~1.25, p = 0.0370). SCZ DEPs exhibited higher enrichment (enrichment ~1.4, p < 0.0001), and SCZ sheddome^14^, i.e. proteins originating from membranes, had the most significant enrichment (enrichment ~2.2, p = 0.0051) (**Extended data Fig.1b**). To select CSF DEPs with a potential role in cognition, we overlapped our SCZ DEPs with published mass spectrometry datasets from AD, BIP, and MDD to generate a “cross-cognitive disorders proteome” (**Supplementary Information Table 3**). This analysis reveals that DEPs shared between several diseases are over-enriched in proteins present in the postsynaptic density (PSD)^29^ (**Fig 1.e, Fig. 1f**). One synaptic DEP was recurrently altered in SCZ and across other brain disorder proteomes: α2δ-1, encoded by the CACNA2D1 gene (pLi >0.9) (**Fig 1.g**). Notably, dysregulation of Cacna2d1/ α2δ-1 occurs cross-species as it is also reduced in the CSF of aged mice^30^. Membrane-bound α2δ-1 is a known target of gabapentinoids and is typically observed at excitatory synapses^29,31^, where it acts primarily as an auxiliary subunit of voltage-gated calcium channels (VGCCs), a gene family consistently associated with NDDs^9,10^. Indeed, rare disruptive mutations in the CACNA2D1-3 genes have been repeatedly found in subjects with SCZ ^32^, and eight protein-truncating variants in CACNA2D1 were recently found in SCZ^9^. Furthermore, complete loss of function of CACNA2D1 leads to a severe, early-onset NDD in humans ^33^. Peptide mapping from our proteomic screen showed that CSF α2δ-1 does not present the canonical transmembrane and intracellular domains, consistent with the shedding of the membrane-associated protein (**Fig. 1h**). We confirmed the 40 % reduction of soluble α2δ-1 protein by western blotting in an independent cohort of individuals <5 years after the initial presentation of psychosis vs. aged/gender-matched unaffected control (**Fig. 1i**). We did not observe any correlation with duration of illness (p = 0.35) or antipsychotic use (p = 0.97) (Table S4). We further replicated the reduced α2δ-1 protein levels in a third independent cohort of post-mortem SCZ CSF samples (**Extended data Fig.1c**). Altogether, we show that the CSF levels of the α2δ-1 ectodomain were consistently reduced in SCZ and broadly across cognitive disorders.

**Fig. 1:**
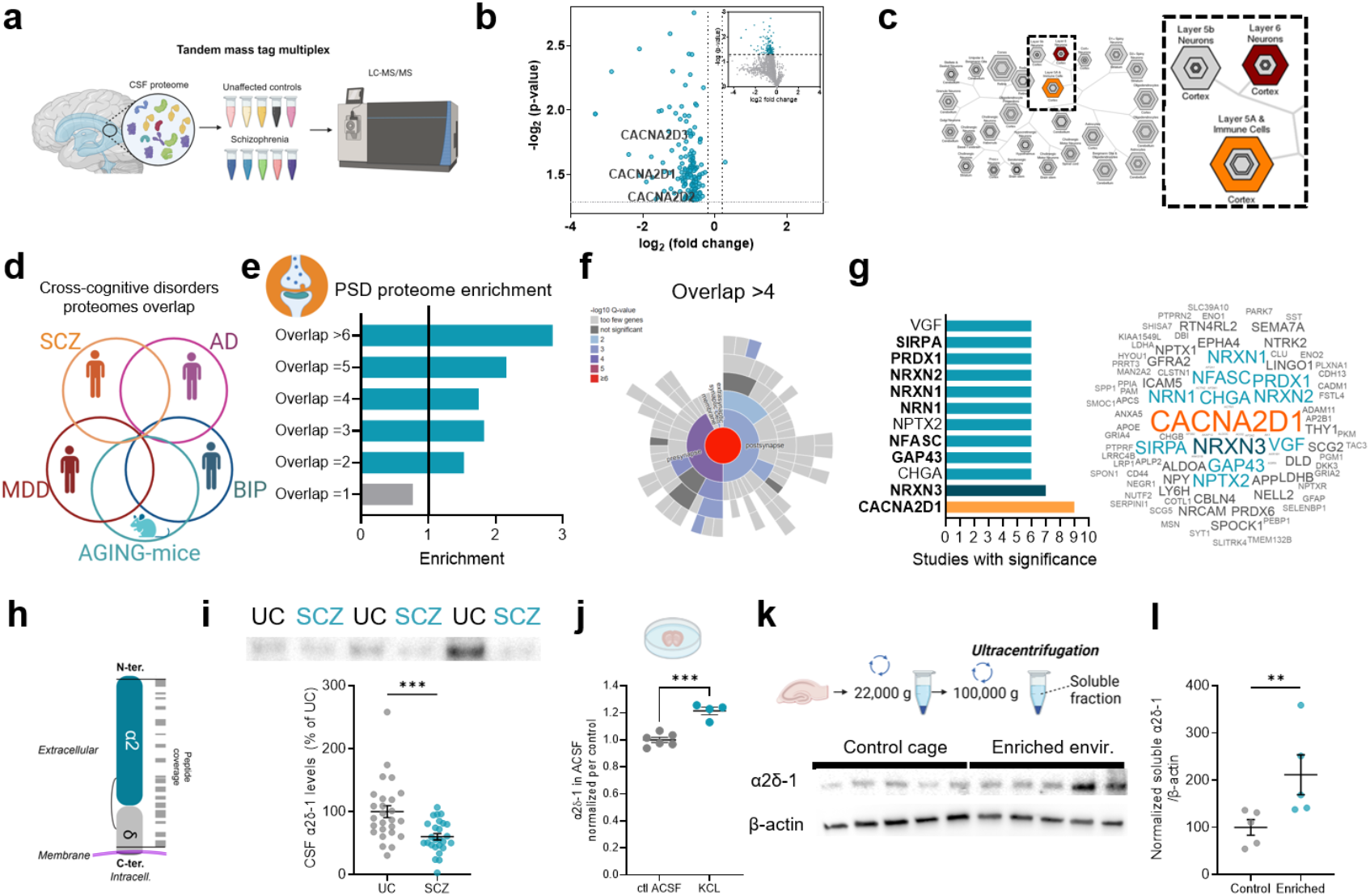
Quantitative proteomics reveals reduced soluble α2δ-1 in CSF of individuals with SCZ. **a**. Experimental schematic of multiplex Tandem Mas Tag proteomics of CSF samples. **b**. Volcano-plot of proteins significantly dysregulated in SCZ CSF *vs*. unaffected controls. **c**. Brain region-specific expression analysis shows enrichment of DEPs only in Layer 5b and 6 cortical genes; gray indicates no statistical significance. **d**. Schematics of the diseases included in different brain disorders proteomes overlap, SCZ: Schizophrenia, AD: Alzheimer’s disease, MDD: depression, BIP: bipolar disorders. **e**. Post Synaptic Density (PSD) proteome enrichment for proteins dysregulated in overlapping proteome datasets, significant over-enrichment is shown in blue (p<0.05). **f**. SynGo analysis of DEPs present in 4 or more datasets. **g**. Left: CACNA2D1, encoding α2δ-1, was identified in 9 out of 9 diseases proteomes, in bold: synaptic proteins. Right, word cloud with the size of the word proportional to their presence in the diseases datasets. **h**. Map of α2δ-1 peptides found in CSF. **i**. Western blot validation of reduced α2δ-1 in an independent cohort of SCZ subjects (<5 years after psychosis, n=26) *vs*. unaffected controls (UC) (n=26, Mann-Whitney test, ***: p<0.001). **j**. Levels of α2δ-1 in the ACSF following high KCl stimulation vs control ACSF, t-test. P<0.05, 4-6 wells with slices from 4 mice. **k**. Methods (Top) and western blot of α2δ-1 (bottom) in the hippocampal soluble fraction of mice in a control cage *vs*. enriched environment. **l**. Quantitation of α2δ-1 normalized to β-actin (n=5 mice/condition, t-test, **: p<0.001).

### Soluble α2δ-1 release in the brain is activity-dependent

We investigated whether the release of α2δ-1 into the extracellular milieu is a regulated, rather than a constitutive, process that might be influenced by neuronal activity or proteolytic enzymes^14^. To examine this possibility, we stimulated acute cortico-striatal brain slices by bath application of either a depolarizing ACSF containing a high KCl concentration (30 mM) or ACSF containing the GABA_A_ receptor antagonist bicuculline (30 μM) for three hours. We then measured α2δ-1 release in the bathing ACSF solution using western blots and found that both stimulation methods increased soluble α2δ-1 levels (**Fig. 1j, Extended data Fig. 2a**,**b**). We next examined this process *in vivo* by exposing mice to either a control cage or an enriched environment for 30 min, and we observed increased α2δ-1 levels in the soluble fraction of hippocampal homogenates in western blots (**Fig. 1k,l**). Prior work has shown that membrane-bound α2δ-1 contains a glycosylphosphatidylinositol (GPI) anchor site^34^, a posttranslational modification that can be hydrolyzed by an enzyme reduced during aging: glycosylphosphatidylinositol-specific phospholipase D1 (GPLD1)^16^. We investigated the involvement of GPLD1 in ectodomain shedding in HEK293 cells expressing α2δ-1 and observed increased levels of soluble extracellular α2δ-1 after stimulation of the cells by the cAMP pathway activator forskolin (FSK, 30 μM) and reduced levels in the presence of the GLPD1 inhibitor 1,10 PNT (**Fig. S2c**,**d**). Together, these results reveal that soluble α2δ-1 release is finely regulated by neuronal and enzymatic activity.

### A synthetic soluble Alpha2Delta-1, SEAD1, preferentially binds interneurons

Previous work has shown that several proteins found in CSF can modulate brain function ^11,12,15^. Thus, to determine if soluble α2δ-1 modulates synaptic plasticity, we designed and produced a recombinant mouse α2δ-1 protein based on the tryptic peptides of soluble α2δ-1 identified in the human CSF samples (**Fig. 1h**) with tags for affinity-purification and immunodetection (Fc: human IgG fragment, and Histidine x6)^35^. For simplicity, we called this protein “SEAD1”, for **S**ynthetic **E**ctodomain of **A**lpha2**D**elta-**1** (**Fig. 2a**). We tested neuronal binding of SEAD1 on cortical cultures and used deconvolved confocal microscopy^36^ to precisely observe SEAD1 fluorescent puncta localizing at the surface of dendrites on the two main types of cortical neurons, pyramidal and parvalbumin-positive (PV+), but significantly more (~40%) on PV+ neurons (**Fig. 2b**). Published single-cell RNA sequencing data from the human neocortex (From Allen Brain Atlas)^37^ (**Extended Data Fig. 3a**) shows that the CACNA2D1 mRNA is predominantly expressed by pyramidal neurons yet we found that it preferentially binds to PV+ dendrites suggesting Pyramidal->PV neuron intercellular signaling. Thus, we investigated the molecular mechanism of SEAD1 preferential binding to PV+ neurons membranes^38^. We found at least two conserved positively charged motifs predicted to interact with perineuronal nets (PNNs) in human and mouse α2δ-1 amino acid sequences^39,40^ (**Extended Data Fig. 3.b**). PNNs are an extracellular matrix densely concentrated around PV+ neurons (**Extended Data Fig. 3c**). We then co-injected SEAD1 in the somatosensory cortex of WT mice, with the enzyme chondroitinase ABC (ChABC) to degrade the PNN on one side of the brain, and BSA in the contralateral hemisphere as a control (**Extended Data Fig. 3c**). Immunolabeling for Fc (**Extended Data Fig. 3d**) revealed that digestion of the PNN by ChABC significantly reduced the binding of SEAD1 around PV+ interneurons *in vivo* (**Fig. 2c, Extended data Fig. 3 c-e**).

**Fig. 2:**
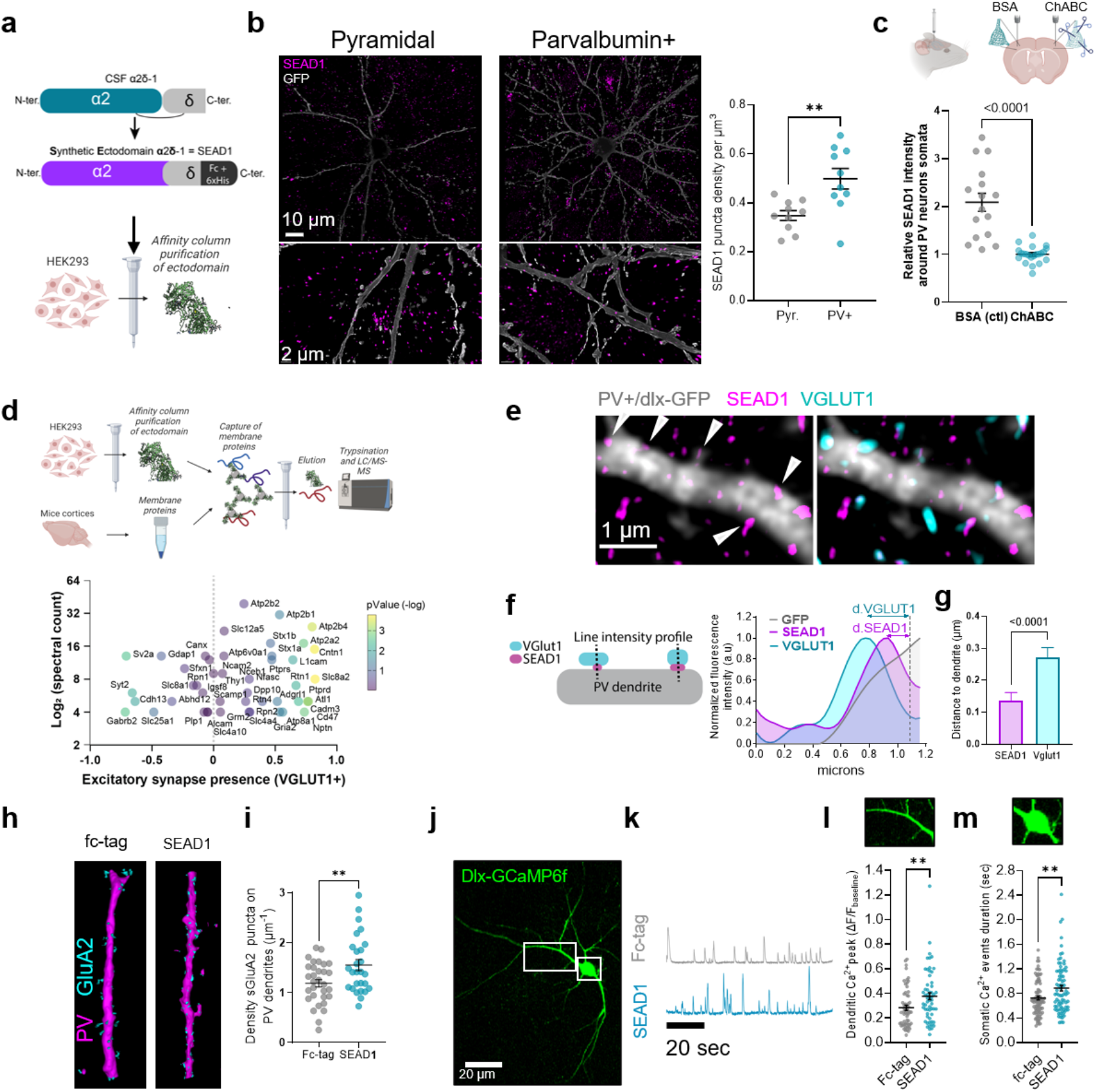
Synthetic Ectodomain of Alpha2Delta-1 (SEAD1) promotes PV+ interneuron dendritic plasticity. **a**. Cartoon schematics of SEAD1 design and purification. **b**. Representative confocal images of SEAD1 binding to cultured GFP-filled (gray) excitatory pyramidal and PV+ neurons and quantification of SEAD1 puncta density/μm^3^ in Pyr *vs*. PV+ neurons (n=10/condition, t-test, **: p<0.01). **c**. Top-Schematic of ChABC enzyme injection to digest the PNN unilaterally in the cortex of adult mice. Bottom: Measurement of average SEAD1 intensity around PV+ neuron somata (n=16-22 neurons from 3 mice, t-test, p<0.0001). **d**. Schematic of SEAD1 target protein affinity pulldown from mouse cortical membrane fraction. **e**. Multiple variable plots of SEAD1 interactors’ enrichment for excitatory synapse presence (1 equal complete correlation with Vglut1 expression), color-coded p-value for correlation with Vglut1 expression. **f**. Schematic and example of normalized line scan intensity profiles for SEAD1, VGLUT1, and GFP immunofluorescence. Distances were measured from the maximum intensity location for SEAD1 or VGLUT1 and GFP (dendrite). **g**. Average distance between VGLUT1/GFP and SEAD1/GFP (n=12, t-test: p<0.0001). **h**. Representative confocal 3D-reconstructed images of surface GluA2 and PV immunofluorescence on dendrites of cultured neurons treated with Fc control or SEAD1 (24 hours). **i**. Quantification of surface GluA2 puncta linear density (n=27-31 neurons/condition, **: p<0.01). **j**. Representative image of Dlx5/6-GCaMP6f neuron. **k**. Representative spontaneous dendritic Ca^2+^ traces from Fc control and SEAD1-treated neurons. **l**. Quantitation of dendritic Ca^2+^ amplitudes (t-test, **: p<0.01, 57-58 dendrites). **m**. SEAD1 increases somatic Ca^2+^ event durations (t-test, **: p<0.01, n=80-88 neurons).

### SEAD1 binds to synaptic protein complexes

We next investigated the molecular targets of SEAD1, which could reflect the protein complexes interacting with soluble α2δ-1. Membrane-bound α2δ-1 has been implicated in regulating synaptic plasticity ^12,15,31,41^, but its membrane protein interactome has not been fully elucidated. Through affinity pulldown from mouse cortical membrane preparations using SEAD1, followed by liquid chromatography-tandem mass spectrometry (LC-MS/MS)^11,35,42^ (**Fig. 2d, Supplementary Information Table 5**), we identified multiple membrane-bound interacting proteins, including neuronal adhesion molecules such as Ptprd, Cadm3, Cntn1, Ncam2, Cadherin-13, L1Cam, and Igsf8; ion channels, Gria2 and Gabrb2, transporters: Kcc2; and the Ca^2+^ export pump Atp2b2 (**Fig. 2d**). Remarkably, SEAD1-interacting proteins were significantly enriched for proteins found the postsynaptic density proteome^29^ and a combined list of genes impacted by *de novo* variants in SCZ^25^, relative to the background of all detected proteins (**Extended data Fig. 4a, Supplementary Information Table 6**). SynGO^43^ analysis showed enrichment for synaptic terms, including presynaptic compartment and integral component of postsynaptic density membranes (**Supplementary Information Table 7, Extended data Fig. 4b**). Notably, a larger number of interactors were enriched in excitatory (VGLUT1+) (**Fig. 2d, Extended Data Fig 4e**) *vs*. inhibitory (VGAT+) synapses by comparing our dataset to the Syndive database ^44^ (**Supplementary Information Table 8**). We validated this preferential binding *in vivo* by injecting SEAD1 in the frontal cortex/anterior cingulate cortex (ACC) of adult WT mice and found higher colocalization with excitatory (Psd95) compared to inhibitory (Gephyrin) postsynaptic markers (**Extended data Fig. 4c-e**).

Together, these results suggest that SEAD1 could primarily function as an extracellular, trans-synaptic bridge at the interface between excitatory presynaptic boutons and inhibitory neuronal dendrites akin to the process observed with neuronal pentraxins^45^. We tested this using structured illumination microscopy (SIM) of SEAD1 and VGLUT1 on dlx-GFP-labelled/PV+ cultured neurons (**Fig. 2e**). We measured the distance between the centre of SEAD1 or VGLUT1+ puncta and PV+ postsynaptic dendrites (**Fig. 2f,g**). We observed SEAD1 binding midway between presynaptic excitatory boutons and dendritic plasma membranes of PV+ cells (**Fig. 2g**).

### SEAD1 enhances interneuron activity

To investigate the functional consequences of SEAD1 neuronal binding, we assessed the number of excitatory postsynapses on PV+ interneurons following incubation of cultured cortical neurons with either SEAD1 or Fc-tag alone. SEAD1 significantly increased the density of surface-expressed GluA2 AMPA receptor subunits (**Fig. 2h,i**): one of the SEAD1 interactors (**Fig. 2d**), and a potential risk factor in SCZ and other NDDs^46,47^. Consistent with an increased number of synapses, intracellular Ca^2+^ imaging of GABAergic interneurons *in vitro* using the genetically encoded Ca^2+^ reporter GCaMP6f under the control of the Dlx promoter (**Fig. 2j,k**) revealed a ~35% increase of dendritic Ca^2+^ event average amplitude (**Fig. 2l**) and a ~20% increase in the duration of the somatic Ca^2+^ events (**Fig. 2m**), in the SEAD1 treated neurons (**Extended data Fig 5a**,**b**). On the contrary, SEAD1 did not affect Ca^2+^ events in astrocytes *in vitro*, confirming a functional effect on Ca^2+^ signalling predominantly in neuronal populations (**Extended data Fig. 6a-c**). Together, these findings demonstrate that SEAD1 binds to synapses and enhances activity in GABAergic interneurons *in vitro*.

### SEAD1 modulates cortical and hippocampal microcircuit activity

GABAergic interneurons are key regulators of neuronal microcircuit dynamics^48–50^, thus, we investigated how SEAD1 affects local network activity and dynamics in cortical neurons cultured on multi-electrode arrays (MEA)^11,51^. We observed a significant reduction in the spontaneous average firing rate per electrode after SEAD1 treatment *vs*. Fc-treated controls (**Fig. 3a,b**). This effect was associated with network dynamics alteration with a decrease in the number of action potentials per network burst (**Fig. 3c**).

**Fig. 3.**
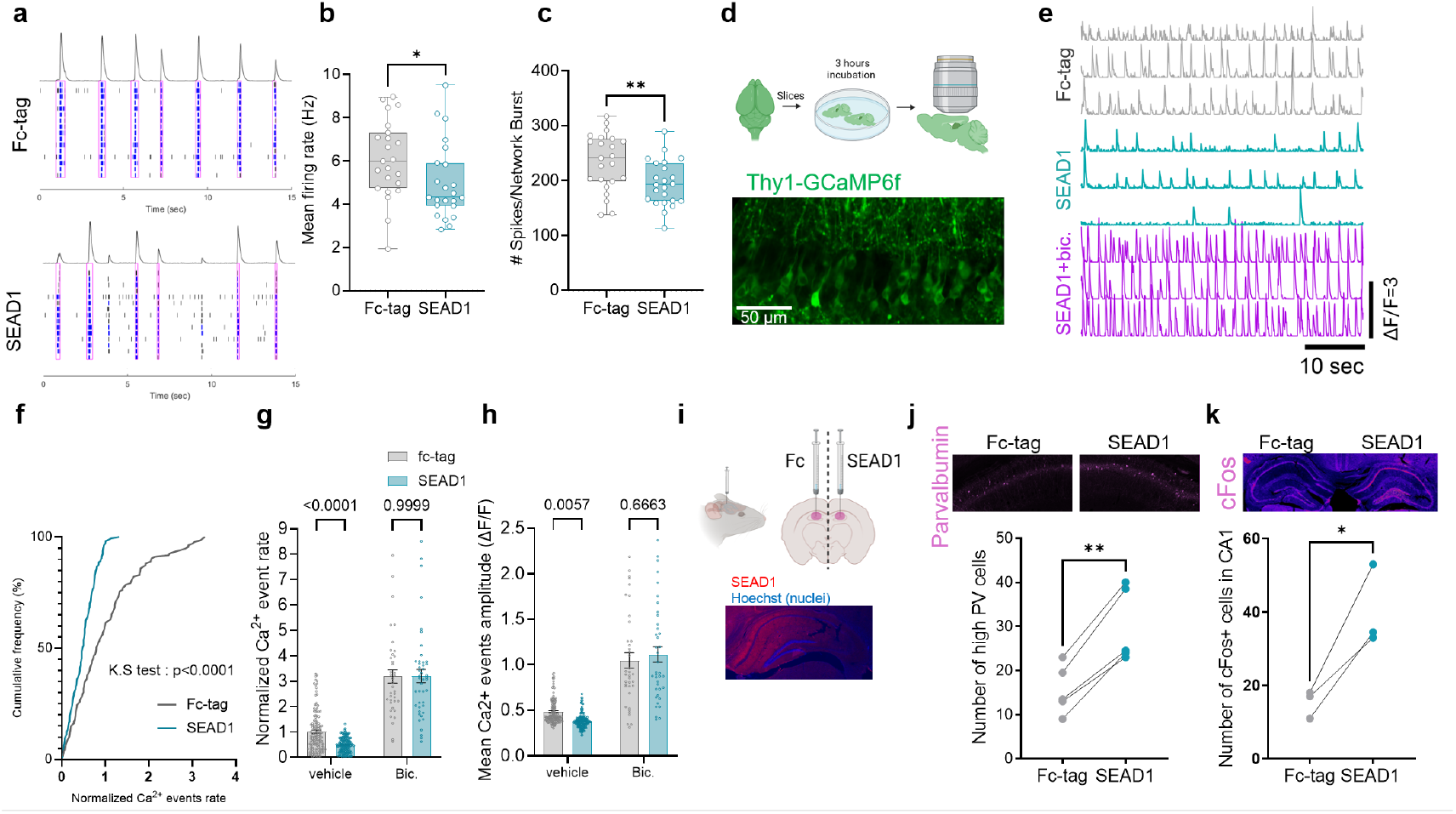
SEAD1 modulates neuronal network dynamics through GABAergic interneuron activity. **a**. Representative raster plots of action potentials detected by MEA in cultured cortical neurons (21 DIV) treated with Fc control or SEAD1 (1 hour). **b**. Quantitation of mean firing rate (n=23 and 23, Mann-Whitney test, p<0.05). **c**. Spontaneous action potentials/network burst (n=23 and 23, t-test, p<0.01). **d**. Schematic and representative image of CA1 hippocampal slices from mice expressing GCaMP6f under the Thy1 promoter. **e**. Representative traces of Ca^2+^ activity in Fc control and SEAD1-treated slices. **f**. Cumulative relative frequency of normalized hippocampal neurons Ca^2+^ event rate (Kolmogorov-Smirnov test, p<0.0001, n=135-154 neurons). **g**. Spontaneous event rates in the absence (n=135-154 neurons, Two-way ANOVA followed by Tukey post-hoc tests) or presence of bicuculline (30 μM) (n=37-45 neurons, t-test, n.s). **h**. Ca^2+^ event amplitudes (Two-way ANOVA followed by Tukey post-hoc tests). **i**. Schematic of hippocampal injection of Fc control or SEAD1 (150 ng) (top) followed by fixation and immunolabeling 6 hours later (bottom). **j**. Representative images of PV immunofluorescence in hippocampal slices from injected mice. Bottom: Number of cells expressing high levels of PV (n=5 mice, paired t-test SEAD1 vs Fc, **: p<0.01). **k**. Number of cFos+ cells on SEAD1 injected side of the hippocampus versus Fc-tag side (n=3 mice, paired t-test, *: p<0.05).

To further validate the impact of SEAD1 on activity in a complete brain microcircuit, we incubated acute hippocampal slices from Thy1-GCaMP6f mice with SEAD1 for three hours before imaging spontaneous network neuronal Ca^2+^ activity in CA1 under a multiphoton microscope (**Fig. 3d,e**). We observed a significant reduction in the spontaneous Ca^2+^ event rate (**Fig. 3f,g**) and amplitude (ΔF/F) (**Fig. 3h**) on the pyramidal neurons somata after SEAD1 treatment, which was completely reversed by the GABA_A_ receptor blocker: bicuculline (30 μM) (**Fig. 3e,g,h**). Thus, both *in vitro* and *ex vivo* data indicate that the stimulatory action of SEAD1 on GABAergic interneurons has an overall inhibitory effect on network activity.

Finally, because PV protein levels are reduced in SCZ and linked to PV+ neuron activity^6^, we asked whether SEAD1 could augment PV expression in WT mice. 6 hours after local *in vivo* injection of SEAD1 into the hippocampal CA1 of adult WT mice (**Fig. 3i,j**), we observed significantly more cells with high levels of PV immunostaining on the SEAD1-injected side *vs*. the contralateral control side (**Fig. 3j**). A similar result was found with the number of cFos+ cells, an immediate early gene typically observed in response to pro-plasticity treatments (**Fig. 3k**). Given that PV and cFos expression are typically increased in the hippocampus during learning^52^, our results suggest that SEAD1 may exert pro-cognitive effects by promoting PV+ interneuron function.

### SEAD1 restores neuroanatomical and cognitive deficits in a genetic mouse model of SCZ

Excitation/inhibition (E/I) imbalance is a hallmark of several brain disorders, including SCZ, and is thought to contribute to impairments in social and cognitive function^27,53–55^. SCZ is characterized by an increased E/I ratio and reduced PV expression levels^27,56^. Notably, our results suggest that SEAD1 has the opposite effect on E/I ratios. Therefore, we hypothesized that the enhancement of inhibitory neuron activity by SEAD1 could restore E/I balance and improve behavioural functions associated with cognitive deficits SCZ^53,57^. SCZ is a highly heritable disorder and copy number variations (CNVs) are among the most penetrant mutations^22^. In particular, the 16p11.2 microduplication increases SCZ risk ~10-fold^9^, and defines the “16p11.2 duplication syndrome” ^21,25,58^. 16p11.2^dup/+^ mice modelling human 16p11.2 duplication exhibit brain alterations characterized by hyperglutamatergic and hypoGABAergic circuits leading to cortical hypersynchrony^25,59^, reduced PV levels in the prefrontal cortex ^60^, and reduced α2δ-1 in the cortical membrane fraction^25^.

We evaluated the effects of SEAD1 *vs*. Fc control injection on frontal cortex anatomy and behaviour in 5-6 months-old 16p11.2^dup/+^ mice (**Fig. 4a**). We observed fewer high PV-expressing PV+ interneurons in the ACC of 16p11.2^dup/+^ *vs*. WT mice (**Fig.4b,c**). Notably, this was restored to WT levels by a single injection of SEAD1 (**Fig. 4b,c**). The ACC is important for sociability^61–63^ and cognitive function^62,64–66^, and is implicated in SCZ^4,67^. We, therefore, carried out a range of behavioural tests to examine social, cognitive, and motor behaviours (**Fig. 4a**). Male 16p11.2^dup/+^ mice displayed significantly less social preference for new mice in the three-chamber test than WT mice, as previously reported^25,59^; this effect was entirely rescued by SEAD1 injection (**Fig. 4d-e**). There was no significant difference between genotypes in female mice (**Fig. 4e**), suggesting a protective effect of the female sex, as observed in other NDD mouse models^68^, and SEAD1 did not affect social behaviour in female mice. Furthermore, novel object recognition performance was significantly lower in pooled male and female 16p11.2^dup/+^ mice *vs*. WT and was rescued by SEAD1 injection (**Fig. 4f**). Finally, SEAD1 did not induce any adverse effects in 16p11.2^dup/+^ unaffected behaviours such as basal locomotion or spontaneous alternation (**Extended data Fig. 7a-e**). Taken together, these data reveal that SEAD1 can rescue changes in PV+ interneurons physiology and behaviour in adult 16p11.2^dup/+^ mice, supporting the idea that treatments based on SEAD1 or other synaptic ectodomains may provide novel therapeutic avenues for SCZ.

**Fig. 4.**
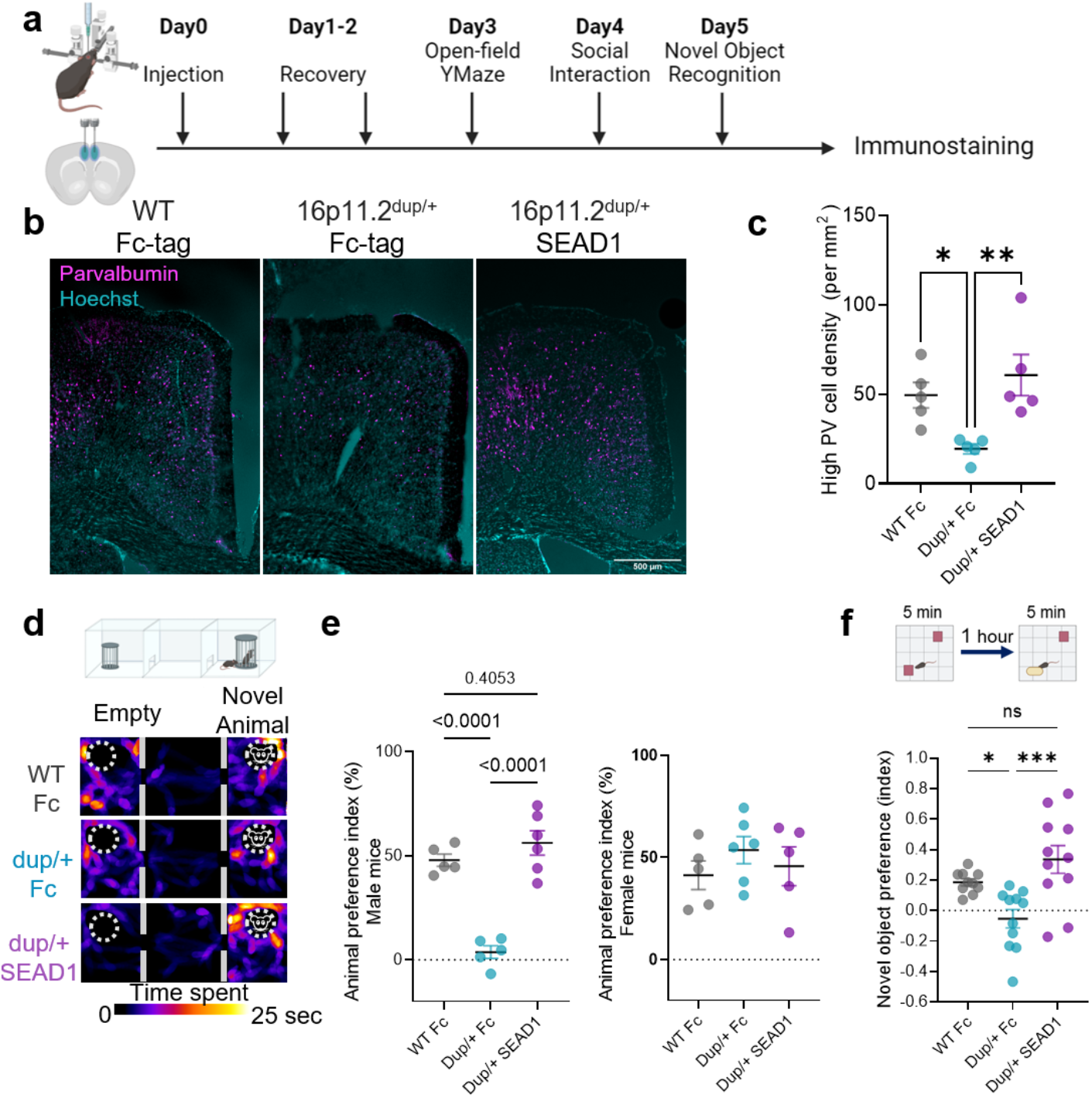
A single SEAD1 brain injection improves anatomical and behavioral endpoints in a mouse model of 16p11.2 duplication syndrome. **a**. Schematic of the experimental design. **b**. Representative images of PV immunolabeling in the ACC of WT, and mutant (16p11.2^dup/+^) mice injected with Fc control or SEAD1. Hoechst labels cell nuclei. **c**. Quantitation of the cell density of PV+ neurons expressing high PV levels (Top 70% cells in fluorescent intensity) (n=5 mice/condition; each point is average from 2-3 slices/mouse; ANOVA and post-hoc Tukey tests, *: p<0.05, **: p<0.01). **d**. Schematic of the three-chamber test (top) and heat map visualization of the time spent by mice in different zones (bottom). **e**. Preference index for stranger mouse *vs*. empty cup in male (left) and female (right) mice upon Fc or SEAD1 injection (ANOVA, post hoc Tukey, p<0.0001). **f**. Schematic of the novel object recognition test (top) and quantitation of the novel object preference index in mice upon Fc or SEAD1 injection (ANOVA, post hoc Tukey, *: p<0.05, ***: p<0.001).

## Discussion

We have shown that the synaptic sheddome detectable in human CSF is altered in individuals with SCZ. In particular, a novel shed form of the α2δ-1 subunit of voltage-gated Ca^2+^ channels is reduced in the CSF in SCZ and several other cognitive decline-related brain disorders. A recombinant soluble α2δ-1 ectodomain, “SEAD1”, acts as a trans-synaptic bridge to modulate pyramidal->PV+ connections, neuronal network dynamics, and rescues behavioural deficits in a genetic mouse model of SCZ. Our work shows that large-scale proteomic analyses can identify global changes in synaptic ectodomain shedding in physiological and pathophysiological conditions, revealing novel mechanisms and biomarkers for disease.

The non-canonical, soluble form of α2δ-1 protein that we have identified in the CSF suggests a potentially novel pro-cognitive mechanism of intercellular signalling that we mimicked using a synthetic protein. Previously, only membrane-bound functions of α2δ proteins were characterized. However, our work reveals that SEAD1 significantly enhances PV+ interneuron function, which reduces neuronal network hyperexcitability.

The interaction of SEAD1 with both pre- and postsynaptic proteins, along with its localization between pre- and postsynaptic sites, suggests a role as a trans-synaptic bridge at excitatory synapses onto PV+ interneurons. Further work could help determine the precise SEAD1 effects on synaptic protein complexes.

Our work also demonstrates the potential of utilizing the CSF proteome to design novel neuronal network modulators. We utilized circuit modulatory properties of SEAD1 to normalize behavioural phenotypes in a mouse model of 16p11.2 duplication syndrome, a prominent risk factor for SCZ, autism, and other NDDs in humans^21,25,58^. The 16p11.2 microduplication region spans 27 protein-coding genes, making gene therapy and drug target identification difficult. Thus, circuit-level alterations may be preferred therapeutic targets for CNV-linked disorders The SEAD1 treatment was remarkably efficient in rescuing phenotypes in adult 16p11.2^dup/+^ mice, an age where antipsychotics targeting dopamine D2 receptors were less efficient in rescuing cognitive deficits in 22q11.1 deletion mice, another CNV mouse model of SCZ^53^. The development of treatments to restore synaptic physiology in SCZ, AD, and MDD remains a high priority, particularly as cognitive symptoms remain largely unaddressed. Consistent with our findings, recent studies have identified proteins in blood plasma^16^ and CSF^15^ that have pro-cognitive properties in aged mice. Because inhibitory neuron dysfunction, disrupted E/I balance, and reduced cognitive flexibility are found in SCZ, ASD, and AD ^54^, soluble α2δ-1-based treatment may represent a potential therapeutic option for a broad range of mental disorders. Recently, α2δ-1 protein levels were found to be reduced in post-mortem brains of COVID-19 patients, further suggesting SEAD1 could have therapeutic potential for the brain impact of this yet untreatable disease. Altogether, our results show that large-scale screens of synaptic ectodomains present in CSF can be directly translated into therapies that normalize the function of brain circuits known to be altered in numerous brain disorders.

## Acknowledgments

The authors would like to thank the Behavioral Phenotyping Core at Northwestern University (Chicago) and its director: Dr. Craig Weiss, for the expertise and equipment. We would like to thank Professor Nicolas Katsanis (Duke University) for providing the backcrossed 16p11.2dup/+ mice. The authors are very grateful to the Nikon Imaging Center, and its director, Dr. Constadina Arvanitis, for the use of the microscopes and technical advice. We also thank Dr. Jones Parker for his helpful comments and suggestions on the manuscript.

## Funding

National Institute of Neurological Disorders and Stroke 5R01NS114977-04

National Institute of Mental Health 5R01MH097216-07

Brain & Behavior Research Foundation (32147)

## Author contributions

MDS and PP conceptualized the entire study. MDS, EBW, and JNS developed the proteomics methodology. MDS, EBW, MB, MF, ELP, SLE, BE, CL, MDMdS, LNS, JMC participated in other experiments methodology. MDS, EBW, MB, MF, ELP, SLE, BE, CL, MDMdS, LNS, JMC participated in the investigation. MDS, KY, EBW, and MF performed data analysis and visualization. Funding acquisition was done by PP and JNS. PP supervised the study. Writing of the original draft was made by MDS and PP.

## Data and materials availability

The CSF proteome and SEAD1 pulldown MS data generated in this study will be deposited in the MassIVE and ProteomeXchange databases after acceptance of the publication. Tools generated in this study will be shared after contacting Peter Penzes.

