## Supplementary material for "Pro-cognitive reshaping of neuronal dynamics by a human CSF-based factor": Methods and supplemental figures

**The PDF file includes:**

Methods
Extended data Fig. 1-7

### Methods

Animal experimentation and care

All animal procedures were performed with the approval of the Institutional Animal Care and Use Committee (IACUC) at Northwestern University. Wild-Type mice experiments were performed on C57bL/6J WT from Jackson Laboratory (Ref 00664). The Thy1-GCaMP6f (C57BL/6J) were purchased from Jackson Laboratory, and heterozygous mice were used for acute slice calcium imaging at 4-5 weeks. 16p11.2 duplication mice were generated by Dr. Alea Mills (Cold Spring Harbor Laboratories). Mice were backcrossed for more than 5 generations to C57BL/6J mice and crossed in our facility with the same strain of female mice. Mice were housed up to 5 animals per cage on a 14h on/10h off light-dark cycle with a temperature of 21-24 ° C and humidity around 40-50%.

Antibodies and pharmacological Reagents

Antibodies were purchased from commercial sources and used as advised by the manufacturer's datasheets: anti-Cacna2d1 (Thermofisher, MA3921) 1/1000 in western blot, anti-Parvalbumin (Thermofisher, PA1-933) 1/500 in IHC and ICC, Anti-gephyrin (Synaptic system 141 011) 1/500 in IHC, Anti PSD95 (Neuromab mouse, K28) 1/500 in IHC, anti-GFP (Abcam chicken, ab13970) 1/2000 in ICC, anti -VGLUT1 (Synaptic System, 135 304) 1/1000 in ICC, anti-Beta actin (Sigma-Aldrich, A5316) mouse 1/2000 in western blot, Anti-human Fc-Cy3 (Sigma C2571) 1/250 in ICC/IHC, anti-human Fc-HRP 1/1000 in Western blot, WFA-FITC: L32481 Thermofisher 3 µg/ml, anti-GluA2 conjugated with Alexa488 for surface staining (1/500) on Neurobasal media for 5 min. Human IgG Fc fragment was used as a control protein (ChromPure Human IgG, Fc Fragment, Jackson Laboratories) for functional and biochemistry experiments. Drugs used were Bicuculline, APV, 1,10 PNT (Tocris).

Plasmids purchased and generated.

pAAV-mDlx-GCaMP6f-Fishell-2 and pAAV-mDlx-GFP were a gift from Dr. Gordon Fishell to Addgene (Addgene plasmid # 83899 ; http://n2t.net/addgene:83899 ; RRID:Addgene_83899) and transfected to neurons in culture at 1 µg/ml of Neurobasal. Cntn1-Fc-His was a gift from Woj Wojtowicz to Addgene (Addgene plasmid #72065; http://n2t.net/addgene: #72065 ; RRID:Addgene_72065). CMV-GFP plasmid was purchased from Origene. We custom-made the plasmid expressing the extracellular domains of Cacna2d1-fc using classic molecular biology techniques. The sequence was extracted from Ensembl to generate the primers. Then, the open reading frame sequence was amplified from the mouse cDNA we generated, and the gel PCR band was cut and then purified using a Takara DNA extraction kit. The insert was then fused to the CMV-GFP plasmid vector. We then used PCR to amplify a truncated form stopping just before the GPI anchor motif to create an endogenously secreted protein. The sequence was then inserted using homologous recombination (Takara Infusion kit) in the vector backbone of Cntn1-fc-his with the CNTN1 open reading frame removed from the plasmid using the restriction enzymes sites: NotI and SpeI.

Human CSF demography

For the Tandem mass tag quantitative proteomics, CSF samples were obtained from Human Brain and spinal fluid Center, Los Angeles (USA), through NIH Neurobiobank, with an average age of 43.2+/-8.5 YO for 4 males and 1 female unaffected controls and 40.8+/5.63YO for SCZ subjects, 3 males and 2 females, no significant difference in age between (+/- standard deviation) groups was observed (t-test, p=0.61).

Replication of the reduction of CSF Alpha2delta-1 was performed on schizophrenia subjects <5 years after the first episode of psychosis (Table S5). These biospecimens were collected from study participants recruited by clinical cohorts established at the Johns Hopkins Schizophrenia Center (*1-3*). This study was conducted with the approval of the Johns Hopkins Medicine Institutional Review Boards. All study participants provided written informed consent. Patients received the Structured Clinical Interview for DSM-IV (SCID-IV) by study team psychiatrists. General linear regression was conducted to calculate adjusted α2δ-1 levels controlling for sex, age, and race. Mann–Whitney U test was conducted to compare the adjusted α2δ-1 levels between SCZ patients and unaffected controls. Spearman correlation analysis was further conducted to evaluate the potential confounding effects of antipsychotic use (chlorpromazine equivalent dose) and duration of illness on α2δ-1 levels in SCZ patients.

Second replication was performed using the ELISA method, from postmortem CSF samples obtained from the University of Miami, Miller School of Medicine, Florida (USA) and Human Brain and Spinal Fluid Center, Los Angeles (USA) through NIH Neurobiobank, the maximum age of death was set to 65 YO to limit confounding effects of aging.

ELISA of CSF samples

Human postmortem CSF samples from 7 healthy controls and 10 SCZ patients (65 Years old and less) with Schizophrenia from NIH Neurobiobank and analyzed for CACNA2D1 using the Human CACNA2D1 sandwich ELISA Kit (Cat # HUFI03408, AssayGenie; Dublin, Ireland) according to the manufacturer's instructions. Human CACNA2D1 was used as a standard. To obtain measurements within the linear range of the standard curve, specimens of human CSF were diluted 100-fold.

Western Blots of CSF samples

A volume of 4.5 µl of CSF was mixed with 4.5 µl of 2X SDS-glycerol-blue Laemmli buffer (Bio-Rad, 2x) and heated at 95 ° Celsius for 5 minutes. The samples were run on a precast 4%-20% gradient gel (Bio-Rad) for 45 minutes at 200 V and then transferred to PVDF membranes (Transblot, Bio-Rad). Membranes were blocked in TBS with 0.1% Tween and 3% BSA for 1 hour before overnight incubation with a primary antibody against Alpha2delta1 (ThermoFisher). Membranes were then washed in TBS-T three times and incubated for 1 hour with a secondary antibody conjugated with horse radish peroxidase (HRP) and washed again three times with TBS-T. The bands were revealed using the Enhanced chemiluminescence (Bio-Rad) method on a Gel imager, and the integrated intensity of the luminescence was measured on the Bio-Rad system.

Recombinant soluble Alpha2delta1-Fc (SEAD1) preparation

Briefly, HEK293 at 70% confluence were transiently transfected with our custom-made plasmid expressing a soluble SEAD1 for 3 days in serum-free media (OptiMem). Cell viability was assessed daily. After three days of expression, cell debris were cleared using centrifugation at 2,000g followed by a 0.2-micron filtering. Clarified ice-cold media was then allowed to flow on a column filled with Nickel beads binding the 6x Histidine tag, washed with 20 mM of Imidazole (pH=8), and eluted with 300 mM of Imidazole. The imidazole was dialyzed against PBS. The solution was then concentrated to 1 µg/µl using a spin column concentrator (Millipore Sigma, Amicon UltraCell 10K). Recombinant protein purity was analyzed on an acrylamide gel (gradient gel 4-20%, Bio-rad), stained with Sypro Ruby or Coomassie. Only preparations presenting a single SEAD1 protein band at around 170 KDa were used for further experiments. The proteins were then aliquoted in small volumes and flash frozen in liquid $N_2$, then stored for less than 6 months at -80C.

Human CSF TMT Mass Spectrometry

-TMT-MS Sample Preparation

TMT-MS sample preparation was performed as previously described (PMID: 33238128). In brief, 200 µg of each CSF sample was used for TMT-MS sample preparation. Proteins were extacted using methanol-chloroform precipitation and the extracted proteins were then resuspended in 6M guanidine in 100 mM *N*-(2-hydroxyethyl)piperazine-*N′*-ethanesulfonic acid (HEPES). The proteins were further processed via the reduction of disulfide bonds with dithiothreitol and alkylation of cysteine residues with iodoacetamide, followed by digestion for 3 h at room temperature (RT) with 1 µg of LysC (Promega) and then further overnight at 37 °C with 2 µg of Trypsin. The digest was then acidified with formic acid and desalted using C18 HyperSep columns (ThermoFisher Scientific). The eluted peptide solution was dried before resuspension in 100 mM HEPES. Micro-BCA assay was subsequently performed to determine the concentration of peptides. 100 µg of peptide from each sample was then used for isobaric labeling. TMT 10-plex labeling was performed on peptide samples according to the manufacturer's instructions (ThermoFisher Scientific). After incubating for 75 min at room temperature, the reaction was quenched with 0.3% (v/v) hydroxylamine. Isobaric labeled samples were then combined 1:1:1:1:1:1:1:1:1:1, desalted with C18 HyperSep columns, and dried. The combined isobaric labeled peptide samples were fractionated into eight fractions using high pH reversed-phase columns (Pierce). Peptide solutions were dried, stored at −80 °C, and reconstituted in liquid chromatography–mass spectrometry (LC–MS) buffer A (5% acetonitrile, 0.125% formic acid) for LC-MS/MS analysis.

-TMT-MS Analysis

TMT-labelled samples were resuspended in 20 µL of buffer A (5% acetonitrile, 0.125% formic acid), and micro-BCA was performed. 3 µg of each fraction was loaded for LC–MS analysis via an auto-sampler with a Thermo EASY nLC 100 UPLC pump onto a vented Pepmap100, 75 µm × 2 cm, nanoViper trap column coupled to a nanoViper analytical column (Thermo Scientific) with a stainless steel emitter tip assembled on the nanospray flex ion source with a spray voltage of 2000 V. Orbitrap Fusion was used to generate MS data. The chromatographic run was performed with a 4 h gradient beginning with 100% buffer A (5% acetonitrile (ACN) and 0.125% formic acid in $H_2O$) and 0% B (99.875% ACN with 0.125% formic acid ) and increased to 7% B over 5 min, then to 25% B over 160 min, 36% B over 40 min, 45% B over 10 min, 95% B over 10 min, and held at 95% B for 15 min before terminating the scan. Multinotch MS3 method was programmed with the following parameters: ion transfer tube temp = 300 °C, easy-IC internal mass calibration, default charge state = 2, and cycle time = 3 s. MS1 detector was set to orbitrap with 60 K resolution, wide quad isolation, mass range = normal, scan range = 300–1800 *m*/*z*, max injection time = 50 ms, AGC target = $6 \times 10^5$, microscans = 1, RF lens = 60%, without source fragmentation, and datatype = positive and centroid. Monoisotopic precursor selection was set to include charge states 2–7 and reject unassigned. Dynamic exclusion was allowed; $n = 1$ exclusion for 60 s with 10 ppm tolerance for high and low. The intensity threshold was set to $5 \times 10^3$. Precursor selection decision = most intense, top speed, 3 s. MS2 settings include isolation window = 0.7, scan range = auto normal, collision energy = 35% CID, scan rate = turbo, max injection time = 50 ms, AGC target = $6 \times 10^5$, and $Q = 0.25$. In MS3, the top 10 precursor peptides selected for analysis were then fragmented using 65% higher-energy collisional dissociation before orbitrap detection. A precursor selection range of 400–1200 *m*/*z* was chosen with mass range tolerance. An exclusion mass width was set to 18 ppm on the low and 5 ppm on the high. Isobaric tag loss exclusion was set to TMT reagent. Additional MS3 settings include an isolation window = 2, orbitrap resolution = 60 K, scan range = 120–500 *m*/*z*, AGC target = $6 \times 10^5$, max injection time = 120 ms, microscans = 1, and datatype = profile.

-TMT-MS Data Analysis and Quantification

Protein identification, TMT quantification, and analysis were performed using Integrated Proteomics Pipeline-IP2 (Integrated Proteomics Applications, Inc., http://www.integratedproteomics.com/). Proteomic results were analyzed with ProLuCID, DTASelect2, Census, and QuantCompare. MS1, MS2, and MS3 spectrum raw files were extracted using RawExtract 1.9.9 software (http://fields.scripps.edu/downloads.php). Pooled spectral files from all eight fractions for each sample were then searched against the Uniprot human protein database (downloaded on 03/25/2014) and matched to sequences using the ProLuCID/SEQUEST algorithm (ProLuCID ver. 3.1) with 50 ppm peptide mass tolerance for precursor ions and 600 ppm for fragment ions. Fully and half-tryptic peptide candidates were included in the search space, all that fell within the mass tolerance window with no miscleavage constraint, assembled, and filtered with DTASelect2 (ver. 2.1.3) through the Integrated Proteomics Pipeline (IP2 v.5.0.1, Integrated Proteomics Applications, Inc., CA, USA). Static modifications at 57.02146 C and 229.1629 K and N-term were included. The target-decoy strategy was used to verify peptide probabilities and false discovery ratios. A minimum peptide length of five was set for the process of each protein identification, and each dataset included a 1%

FDR rate at the protein level based on the target-decoy strategy. Isobaric labeling analysis was established with Census 2, as previously described. TMT channels were normalized by dividing it over the sum of all channels. No intensity threshold was applied. The fold change was then calculated as the mean of the experimental group standardized values, and *p*-values were then calculated by Student's *t*-test.

Membrane Pulldown and Mass Spectrometry

The neocortices from 8 mice were rapidly dissected on ice and placed in 20 ml of ice-cold homogenization solution containing Hepes 5 mM, NaCl 150 mM, NaHPO4 20 mM, 5% sucrose at pH 7.4 and mechanically homogenized. The nuclei and larger debris were precipitated at the bottom of a centrifuge tube by performing a first centrifugation at 1,500 g. The supernatant was then centrifuged at 22,000 g to pellet the membranes. The pellets were then resuspended in 22 ml of cold homogenization buffer supplemented with Triton (1%) for 1 hour. The solution was then spun at 160,000 g for 2 hours to precipitate the non-soluble proteins. The supernatant containing Triton-soluble membrane proteins was then split into two 15 ml conical tubes and incubated with 250 µg of Agarose A/G beads coated with 100 µg of SEAD1 or Fc fragment alone as a control overnight. The proteins bound to the beads were then eluted in Glycine pH=2 solution, and subjected to Trichloroacetic Acid (TCA) precipitation. In brief, the volume of the elution was adjusted to 400µL with 100 µM Tris-Cl pH 7.5, and the TCA was added at a final concentration 20% (v/v). The samples were vortex-ed, kept in an ice bucket at 4°C overnight, and the next day, they were span-down at 13,000 rpm, 4°C for 30min. The TCA was carefully removed, and the pellets were washed three times with 100% ice-cold methanol and dried at 95°C for 5min.

The precipitated samples were solubilized and denatured in 8M urea for 30min and processed with 0.2% ProteaseMAX (Promega V2072) for 2h. Subsequently, the samples were reduced with 5mM Tris(2-carboxyethyl) phosphine (TCEP) at room temperature (RT) and alkylated with 10mM iodoacetamide (IAA) while protected from light. Then, they were diluted with 50mM ammonium bicarbonate, quenched with 25mM TCEP and digested overnight at 37°C with 1µg sequencing-grade trypsin (Promega V5280). The digestion was terminated using 1% formic acid and the peptides were desalted using Pierce C18 spin columns.

3 µg of each sample was auto-sampler loaded with a Thermo Fisher EASY nLC 1000 or nLC 1200 UPLC pump onto a vented Acclaim Pepmap 100, 75 µm by 2 cm, nanoViper trap column coupled to a nanoViper analytical column (Thermo Fisher 164570, 3 µm, 100 Å, C18, 0.075 mm, 500 mm) with a stainless steel emitter tip assembled on the Nanospray Flex Ion Source with a spray voltage of 2,000 V. Buffer A contained 94.875% H2O with 5% ACN and 0.125% formic acid (FA), and buffer B contained 99.875% ACN with 0.125% FA. The chromatographic run was 4 h in total with the following buffer B profile: 0%–7% for 7 min, 10% for 6 min, 25% for 160 min, 33% for 40 min, 50% for 7 min, 95% for 5 min, and 95% again for 15 min. Additional MS parameters include: ion transfer tube temp = 300°C, Easy-IC internal mass calibration, default charge state = 2, and cycle time = 3 s. The detector type was set to Orbitrap, with 60 K resolution, wide quad isolation, mass range = normal, scan range = 300–1500 m/z, max injection time = 50 ms, AGC target = 200,000, microscans = 1, S-lens RF level = 60, without source fragmentation, and datatype = positive and centroid. MIPS

was set to "n" and included charge states = 2–6 (reject unassigned). Dynamic exclusion was enabled, with n = 1 for 30 and 45 s exclusion duration at 10 ppm for high and low, respectively. Precursor selection decision = most intense, top 20, isolation window = 1.6, scan range = auto normal, first mass = 110, collision energy 30%, CID, Detector type = ion trap, Orbitrap resolution = 30K, IT scan rate = rapid, max injection time = 75 ms, AGCtarget = 10,000, Q = 0.25, inject ions for all available parallelizable time. Raw spectrum files were extracted into MS1 and MS2 files using the in-house program RawXtractor or RawConverter (http://fields.scripps.edu/downloads.php) (He et al., 2015) and the tandem mass spectra were searched against UniProt's mouse protein database (downloaded 09/28/2019) and matched to sequences using the ProLuCID/SEQUEST algorithm (ProLuCID version 3.1) (Eng et al., 1994) with 50 ppm peptide mass tolerance for precursor ions and 600 ppm for fragment ions. ProLuCID searches included all fully and half-tryptic peptide candidates that fell within the mass tolerance window and had with unlimited mis-cleavages. Carbamidomethylation (+57.02146 Da) of cysteine was considered a static modification. False discovery rate (FDR) was set to 1% at the protein level, for all experiments. Peptide probabilities and FDR were calculated based on a target/decoy database containing the reversed sequences of all the proteins appended to the target database (*4*).

Criteria for SEAD1 membrane interactors selection

All affinity-precipitated proteins from the immunoaffinity assay presenting at least 1 spectral count in the Fc (control) were discarded from the analysis. The rest of the potential interactors were then filtered to select candidates presenting at least 3 different unique peptides and at least 4 spectral counts. To select membrane interactors reflecting the extracellular binding of SEAD1, the overlap between interactors and membrane and GPi-anchored proteins from UniProt was performed to create a final list of high-confidence membrane-bound binding partners. Finally, to measure the affinity for excitatory synapse proteins, the overlap between the list of membrane interactors and the PSD proteome was quantified and statistically tested using the hypergeometric test (as in Forrest et al. 2022).

Proteomics bioinformatic analysis

Protein lists from LC-MS experiments were converted into gene name lists for comparison and overlap with the lists present in databases. Sheddome overlap method was adapted from De-Saavedra et al., a list of proteins with a transmembrane domain or a GPI anchor was exported from Uniprot database (uniprot.org) and overlapped with proteomics gene lists.

CSF protein lists from the TMT proteomics were exported to the CSEA tool to find their putative brain origins based on gene enrichment using the CSEA tool (http://doughertytools.wustl.edu/CSEAtool.html). Synaptic Gene ontology (SynGO 1.2, https://www.syngoportal.org/ ) analysis enrichment was performed on CSF and SEAD1 pulldown datasets with the background of geneset represented by the total list of detected proteins in their respective proteomics experiments.

For post synaptic proteome enrichment, we used the list of proteins from Bayes et al (https://www.genes2cognition.org/proteomics/).

"Syndive" (*5*) Single synapse proteomics for Excitatory vs Inhibitory synapse enrichment, list of genes filtered with presence in the Syndive dataset, 48/71 genes are present in both membrane pulldown and van Oostrum et al. 2023. The correlation with VGLUT1 or VGAT was plotted and compared statistically to indicate the enrichment for each synapse type.
Word cloud was generated using the built-in function in Matlab.

Rat neuronal cultures preparation
Rat cortical cultures were prepared from E18 Sprague-Dawley embryos as in (*6*). Briefly, embryos cortices were dissected, and their meninges were gently removed. Then, the cortices were enzymatically dissociated on Trypsin for 15 min and mechanically dissociated. Cells are then passed on a cell strainer to get remove aggregated cells. Cells were then plated on Neurobasal media (+B27, Glutamax, and Penicillin/Streptomycin). Cells were then fed twice a week with half of the well volume replaced with fresh Neurobasal. Cultures were used for experiments from P17-P21.

Confocal and SIM Microscopy:
Multichannel structured illumination microscopy (SIM) images were acquired using a Nikon Structured Illumination super-resolution microscope using a 100× 1.4 NA objective and reconstructed using Nikon Elements software. Cultured rat neurons were sparsely transfected at Div 18 with a plasmid expressing GFP under the control of the Dlx promoter targeting GABAergic interneurons (*7*). Neurons were then incubated for 30 minutes with 50 nM of SEAD1 in aCSF. The cultures were then surface immunolabeled using antibodies anti-PV, -GFP, -VGLUT1, -human Fc fragment conjugated with Cy3. Distances between SEAD1, GFP, and VGLUT1 were measured using line scans. Conventional confocal images were acquired conventional confocal images were performed on a Nikon Confocal C2 scanner using a 60X oil objective, numerical aperture of 1.4, zoom 2, pixel size of 100 nm, and Zstep of 200 nm.

3D image processing and automated image analysis:
Confocal images were deconvolved using automatic 3D deconvolution from the NIS-elements (Nikon) software. SIM images were reconstructed using the automatic 3D SIM reconstruction from NIS elements. 3D puncta of alpha2delta1, surface GluA2, PSD95, or gephyrin were all automatically detected and segmented using the iterative segmentation methods (ImageJ) and then analyzed using the plugin Distance Analysis (DiAna). Objects physically touching were considered as colocalizing.
For analysis of the GluA2 surface staining on PV+ neurons, dendrites were first smoothed using a median filter set at 4 pixels, and dendrite contours were segmented using a threshold based on their fluorescence intensity (95% percentile). GluA2 puncta were automatically segmented in 3D using the "3D iterative Thresholding" plugin and then analyzed using the plugin Distance Analysis (DiAna). Objects physically touching were considered as colocalizing.

In vitro imaging of calcium transients in soma and proximal dendrites in cultures
Inhibitory neurons from Div21 rat cortical cultures were transfected using a plasmid expressing GCaMP6f under the control of Dlx promoter (1 µg/ml of plasmid and

Lipofectamine 2000). The cultures were incubated with 50 nM of SEAD1 or fc-fragment for 48 hours in a cell incubator. Coverslips were then transferred to an imaging chamber in 5% O2 at 37°C and imaged in wide-field fluorescence, illuminated at 488 nm, using a 20x objective, recorded at 20 Hz on a Zyla camera to precisely detect fast $Ca^{2+}$ transients. Each field of view was acquired for 2 min.

For analysis of $Ca^{2+}$ in astrocytes, neuronal cultures at Div21 were incubated with Cal520AM and SR101 for 15 min, washed with ACSF, and imaged on the same microscope described above.

Multi-Electrode Array

Rat neurons were prepared as described above and plated on 48-well multi-electrode array (MEA) plates (Axion). At Div21, neurons were incubated with 50 nM of SEAD1 or Fc for 1 hour before recording spontaneous extracellular action potentials. Neuronal activity was automatically analyzed from action potentials selected as peaks above 6 times the standard deviation of the signal. Wells were discarded if they did not present a mean firing rate of at least 0.1 Hz. The firing rate, synchrony index, and network bursts were measured and statistically tested on Axion software using default parameters.

Intracerebral injections of recombinant proteins

To measure the impact of perineuronal nets for SEAD1 binding, 0.5 µg of the enzyme Chondroitinase ABC (Proteintech, 6877-GH-020) was injected into one side of the S1 cortex of WT mice (C57Bl6, Jackson Lab) with 0.5 µg of SEAD1, the contralateral side was injected with BSA as a control protein and SEAD1. Mice received Meloxicam as an analgesic (20 mg/kg, subcutaneous) and were allowed to recover for 3 hours after surgery before being injected intraperitoneally with Euthasol (50 mg/kg) and transcardiacally perfused with 4% PFA in PBS.

To measure binding between SEAD1 and synaptic markers in vivo (PSD95 and Gephyrin), we followed the same surgical procedure except that the injection site was located in the anterior cingulate cortex (from Bregma, AP: 1.3 mm, Lateral: 0.3 mm, D-V: 1.5 mm). The fixation was also modified to improve PSD95 and gephyrin immunostaining quality by using a solution of Glyoxal 3%, and acetic acid 0.8% at pH=4.5. Brains were post-fixed overnight in the same fixative solutions, PFA, or glyoxal, and then sliced on a Cryostat (Leica) at 60 microns in frozen Embedding Medium (O.C.T. Compound, Tissue-Tek). The same injection site in the ACC was used for the treatment of 16p11.2 dup mice. 5-6-month-old 16p11.2 dup and WT mice were allowed to recover for 48 hours after injection before starting behavioral assays.

Acute slices preparation

For the analysis of the activity-dependent release of endogenous soluble Alpha2/delta-1 in the extracellular milieu, brains from 4 WT mice per experiment aged from 4 to 5 weeks were rapidly dissected and placed in an ice-cold “cutting” Hepes-aCSF for 2 minutes (NaCl 92, KCl 2.5, $NaH_2PO_4$ 1.2, $NaHCO_3$ 30, HEPES 20, glucose 25, sodium ascorbate 5, sodium pyruvate 3, $MgSO_4.7H_2O$ 10, $CaCl_2.2H_2O$ 0.5, pH 7.3 and bubbled with 95% $O_2$, 5% $CO_2$). The brains were cut on their base at the level of the cerebellum and then glued on a vibratome stage (Leica VT1000s), cut at speed: 1, vibration speed:

8, 250/300 µm thickness. Coronal slices were then placed in the Hepes-aCSF at 32 ° C for 1 hour before starting the stimulation. Two slices per well were then placed in control aCSF NaCl 124, KCl 2.5, $NaH_2PO_4$ 1.2, $NaHCO_3$ 24, HEPES 5, glucose 12.5, $MgSO_4.7H_2O$ 1, $CaCl_2.2H_2O$ 2, pH 7.3, 95% $O_2$, 5% $CO_2$ in a plastic well with 700 µl of aCSF with either vehicle (DMSO or water) and KCl (30 mM) or Bicuculline 30 µM) as neuronal activity enhancers. Slices were left for 3 hours before the aCSF was gently aspirated and placed in a microcentrifuge tube. The aCSF was then cleared by centrifugation at 1,500 g, then 100,000 g for 2 hours. The soluble supernatant was collected, and protein concentration was quantified using the BCA assay (Biorad). 2 µg of proteins were used per lane of the acrylamide gel used for subsequent western blot.

Multiphoton imaging of *Ex Vivo* imaging of hippocampal activity:
Slices from 4-5 weeks Thy1-GCaMP6f (Jackson Laboratories) mice were prepared using the same method presented above, except that after the recovery period, half of the slices were incubated with 50 nM of SEAD1 or Fc fragment for at least 3 hours at 32°C before being placed on a live imaging stage (Warner Instrument). Slices were stabilized using a stainless-steel harp (Warner) with nylon threads. A peristaltic pump was set to superfuse 32°C aCSF to the brain slice with a flow of 4-5 ml per minute to maintain correct oxygenation. aCSF was continuously aerated with a 95% O2, and 5% CO2 gas mixture, and the imaging chamber was also heated at 32 ° Celsius.
Neuronal calcium transients were acquired in the CA1 region of the dorsal hippocampus, at a depth of around 30 µm under the surface of the slice on a Multiphoton Leica (25x, NA = 0.95, Leica), using the resonant scanner to image at 10 Hz on the Hybrid detector (Leica). The femtosecond pulse laser (Coherent) was set at 920 nm, with a power of around 20-30 mW in samples. In these illumination conditions, no more than 5% photobleaching or any visible blebbing during recording was observed in any of the slices. Some slices were imaged with Bicuculline (30 µM) in the bath to inactivate fast inhibitory signaling.

Calcium peaks detection and synchrony analysis:
A z-score map was created from calcium imaging movies on ImageJ to segment ROIs corresponding to neuronal soma with a maximum ZScore>2.5 for the duration of the recording to select active neurons. The contour of the cells was then exported to the ImageJ ROI Manager, and the mean intensity for each frame was then calculated and analyzed on Matlab.
Synchrony analysis: Calcium activity traces for each neuron were normalized in Matlab by subtracting the raw fluorescence signal (F) with the baseline defined as a moving median window of 20 seconds (F0) and divided by the baseline: (F-F0)/F0, or $\Delta F/F$). Calcium peaks were then detected automatically using the “FindPeaks” Matlab built-in function with a minimum of 15% fluorescence variation above the baseline. Peak amplitude and frequency were extracted from the outputs of the function. We then determined synchronized events by performing 10,000 random permutations of the calcium events for each neuron and calculated the minimal number of co-active neurons considered as a network event equal to the 99th percentile of the sum of events for the permutated dataset. We then calculated the percentage of co-active cells during each of these synchrony frames and averaged them for each experiment.

Behavioral Assays:

To measure the *in vivo* impact of brain activity on soluble Alpha2delta-1 cleavage, mice were placed in an ~80 cm square open field in the presence of mouse toys (tunnels, wheels, cubes, houses) for 30 minutes and were then rapidly anesthetized with 5% isoflurane, and their brains were dissected and flash-frozen in liquid $N_2$. The soluble fraction was prepared from the whole hippocampal lobes using sequential centrifugation, as explained above for the membrane pulldown.

16p11.2dup mice and WT littermates were injected in their ACC on day0 with 150 ng of recombinant SEAD1 in sterile PBS and allowed to recover for two days in their home cage with daily meloxicam injections (20 mg/kg) as analgesics.

Before starting the behavioral analyses on 16p11.2 dup mice compared to WT, animals were briefly habituated on day2 with the Open-field (a grey opaque square plastic box of around 80x80 cm), Y-Maze, and the three chambers arena for 20 min per apparatus. All the experiments are carried out and analyzed blind to the genotypes and treatments. Then on day3, animals were individually placed on the Open field in a soundproof room and imaged from the top with a 16-fps gray-scaled camera movies were then further analyzed on ImageJ by tracking the centroid of the animals for each frame and summing the total distance traveled. They were tested the same day for spontaneous alternation, animals are placed in a 3-arm Y maze and let freely explore the apparatus for 5 min. On day 4, animals were tested for sociability in a three-chamber arena. Briefly, animals were first habituated with the arena with two empty cups placed equidistant from the most lateral chamber (see figure 5.D). A stranger conspecific, same sex, was then placed under one of the two wired cups and the mouse tested was let to freely explore again for 5 min. The amount of time each mouse spent investigating the empty cup or novel mouse (defined as sniffing or interacting) was manually recorded. The data for each mouse was converted to a discrimination index (DI (%) = $[(t_{mouse}-t_{object})/(t_{mouse}+t_{object})\times 100]$). On day 5, animals were tested for novel object recognition, first, animals were placed in the open-field arena with no object for 5 minutes. They were then exposed to two identical objects placed on opposite corners of the arena for 5 min. Animals are removed from the arena and allowed to rest in their home cage for 1 hour before reexposing them to the arena with one of the two objects replaced by a new object with approximately the same dimensions. The amount of time each mouse spent investigating the familiar object or novel object for each mouse was converted to a discrimination index (DI = $[(t_{Novel}-t_{Familiar})/(t_{Novel}+t_{Familiar})]$).

Statistical analysis:

All statistical tests were performed using GraphPad Prism, R, or Matlab. Experiments were performed blind to the conditions, and for animal rescue experiments, experimenters were blind to both genotypes and treatments. Bar graph depicts mean +/- SEM except if specified. Significance was defined at $p<0.05$ in all the statistical tests performed in this study. The normality of the data distribution was tested using the Shapiro-Wilk normality test. For tests with two conditions, a t-test was used if data were normal, otherwise, non-parametric test: Kolmogorov-Smirnov or Mann-Whitney tests were used. ANOVA and Two-way ANOVA followed by Tukey post-hoc were used for experiments with more than two conditions. Hypergeometric test for gene set analysis were performed in R. Synchronized networks frames on calcium imaging videos were performed using custom-built scripts on Matlab adapted from (https://github.com/marcdossantosPHD/PenzesLabimaging).

### Extended data figures

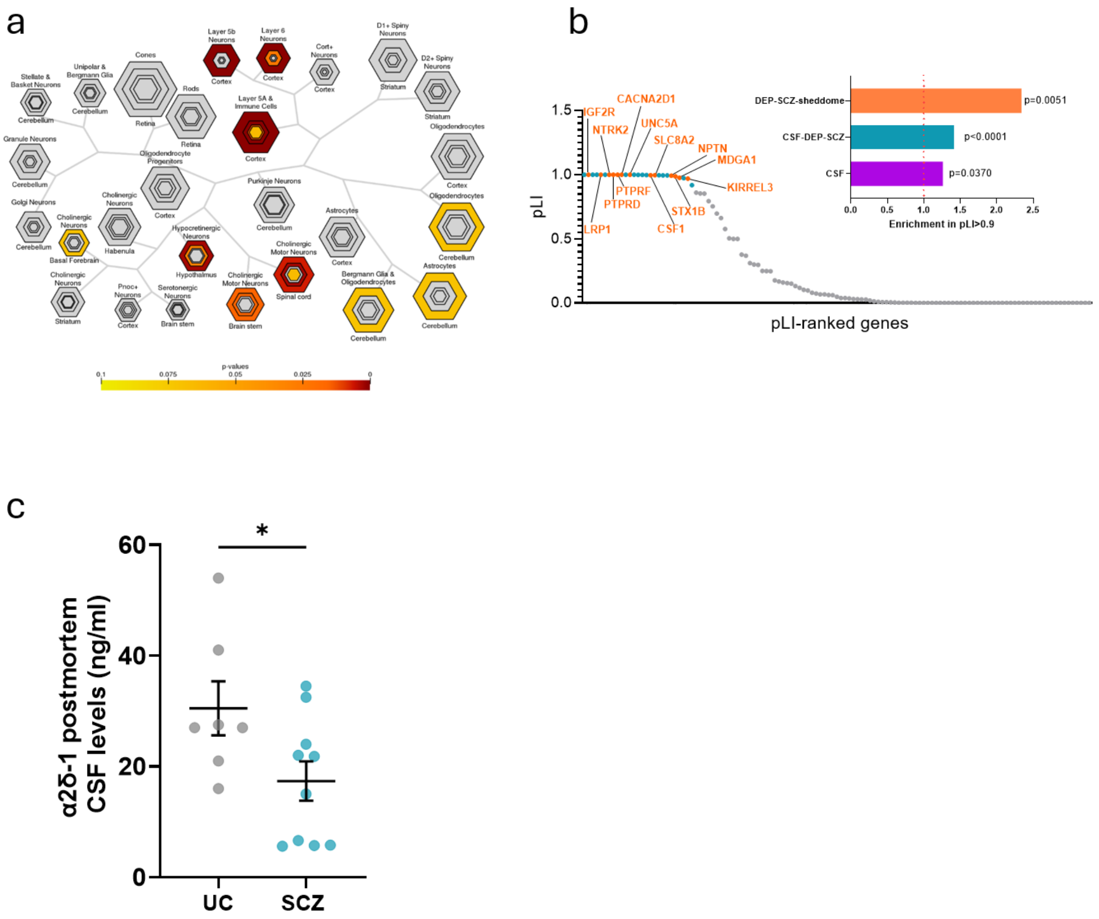


**Extended data Fig. 1: Total proteome in CSF putative brain origin is widespread.**
**a.** Brain region Specific Expression Analysis diagram showed enrichment in Layer 5b and 6 cortical genes when using the total detected CSF protein. The gray color indicates no statistical significance. **b.** Plot of pLi per DEPs; in blue: DEP pLI>0.9, in orange DEPs with pLi>0.9 and a transmembrane domain. **c.** ELISA validation of reduced α2δ-1 in a second cohort of post-mortem CSF samples from SCZ (n=11) *vs.* matched controls (UC, n=7) (unpaired t-test, *: $p<0.05$).

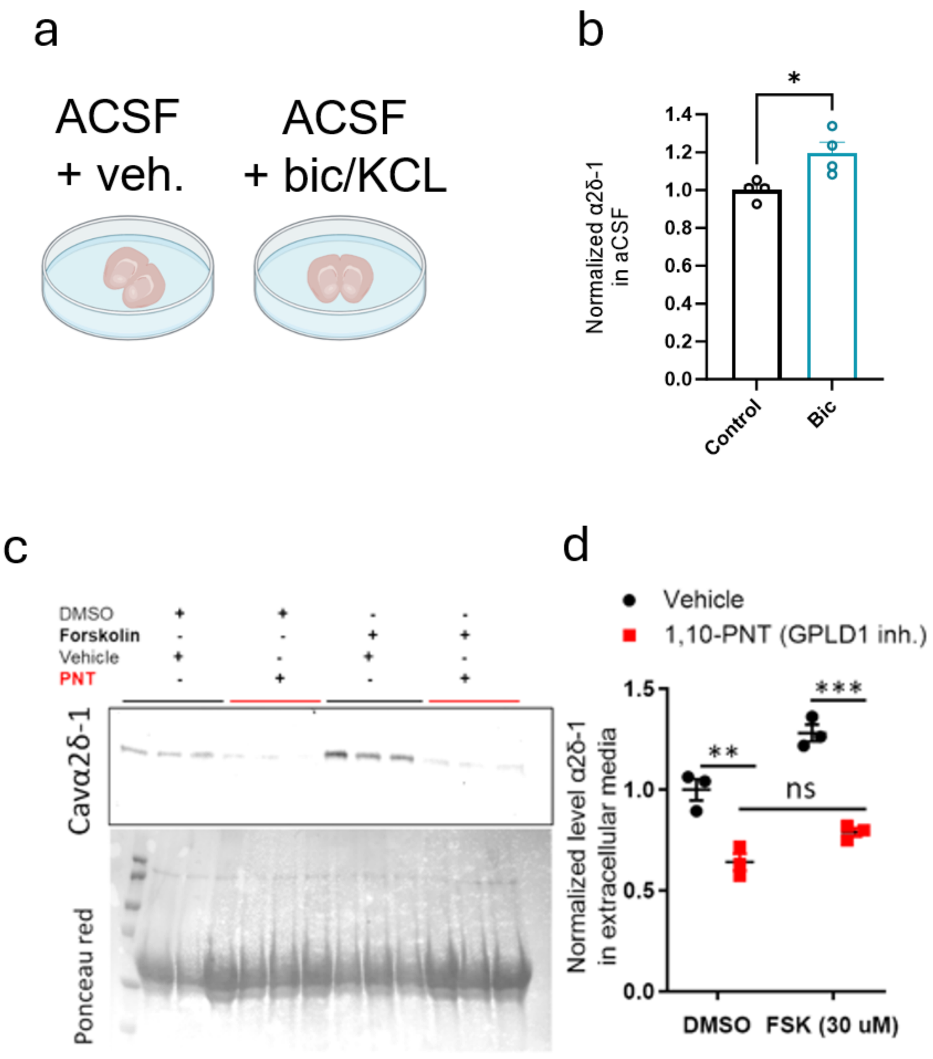


**Extended data Fig. 2: Cellular and molecular regulation of endogenous α2δ-1 release in acute brain slices and in vitro in cell lines.**
**a.** Schematics of the brain slices used for α2δ-1 ex vivo secretion analysis **b.** Levels of α2δ-1 in the aCSF following Bicuculline (Bic, 30 μM) stimulation vs control ACSF, t-test. P<0.05, 4 mice. **c.** Hek293 cells were transiently transfected with a plasmid expressing mouse membrane-bound α2δ-1 and treated with cAMP agonist, Forskolin (30 μM) or a GPLD1 inhibitor 1,10 PNT. **d.** Quantitation of the α2δ-1 blots, relative to DMSO-vehicle (two-way ANOVA, post-hoc Tukey, 3 wells per condition).

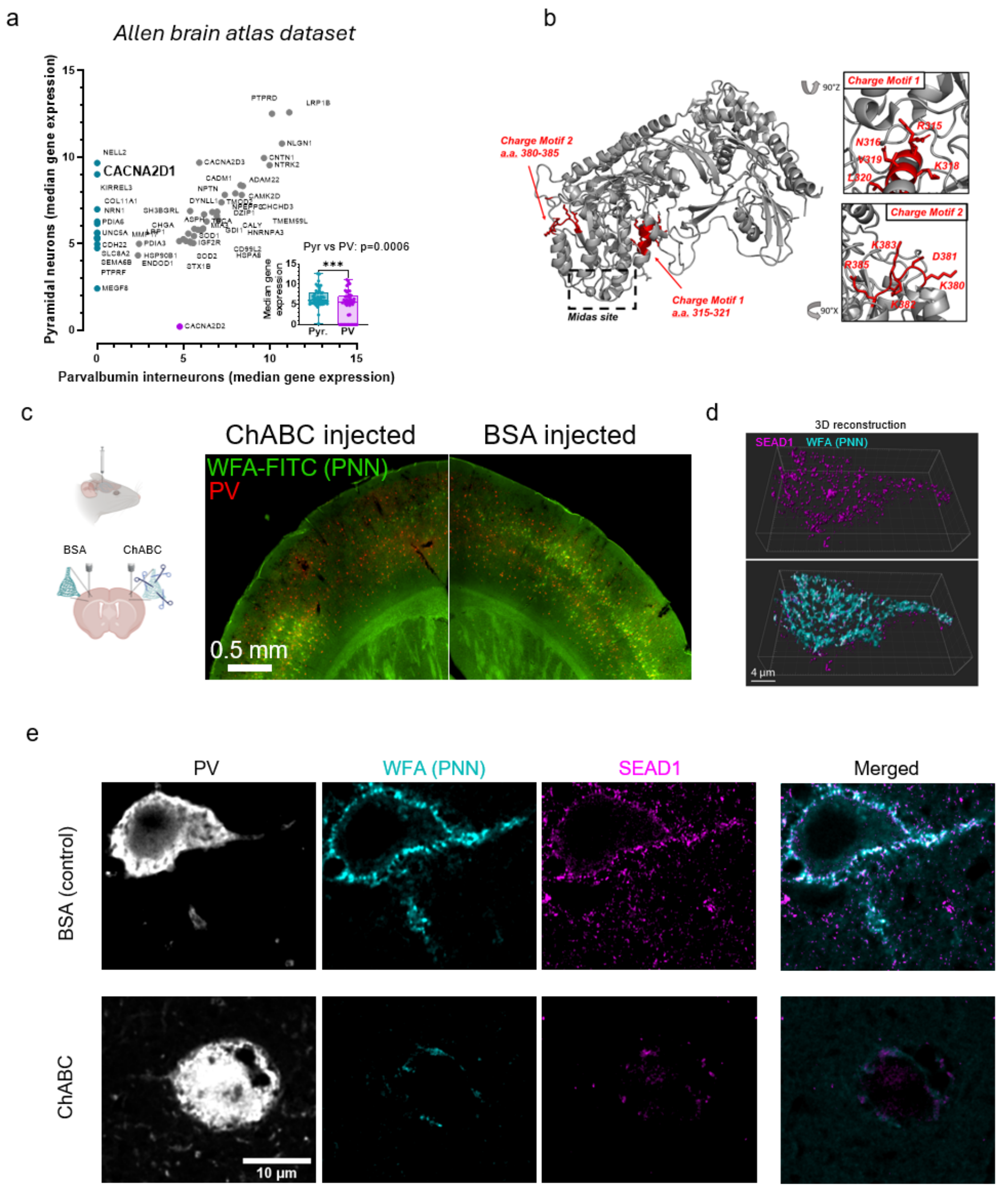


**Extended data Fig. 3. SEAD1 has a high affinity for PNN-surrounded cells in the cortex *in vivo*.**

**a**. Graphical representation of CSF gene expression in excitatory neurons and PV interneurons from single-cell RNA sequencing data. **b**. 3D representation of the 2 amino acid motifs on α2δ-1 predicted to have the charge motif predicted to increase perineuronal nets binding. **c**. Left: cartoon of intracranial injection of chondroitinase ABC (ChABC) or BSA in mice**.** Right: representative image of the PNN stained with WFA-FiTC (green) digestion using ChABC without affecting PV expression (Red). **d**. 3D representative image of high-magnification confocal microscopy of SEAD1 binding PNN positive cells. **e**. Representative images of single confocal microscopy planes showing reduced SEAD1 staining around PV neurons after digestion by ChABC.

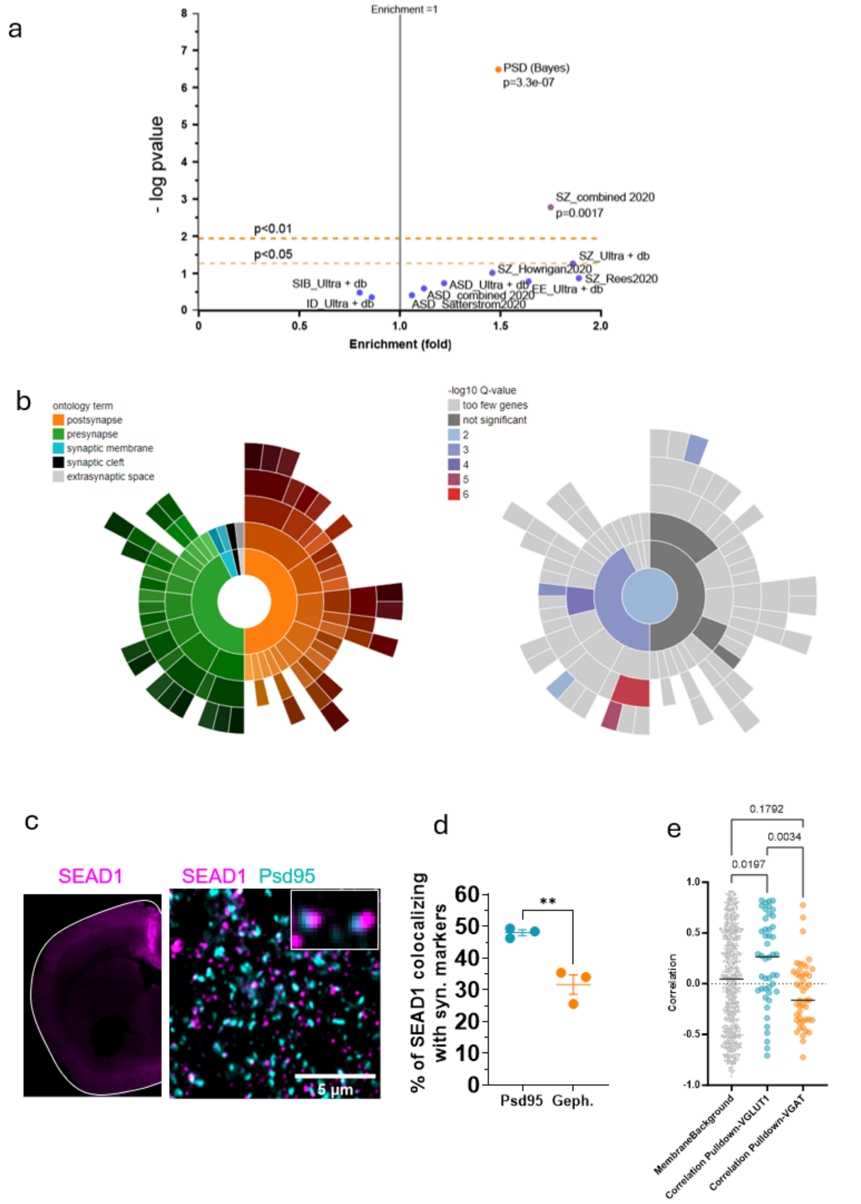


**Extended data Fig. 4. Enrichment for excitatory synaptic proteins and schizophrenia risk genes among the SEAD1 interactors.**

**a.** SEAD1 interactors are over-enriched for the PSD proteome as well as combined *de novo* mutants for schizophrenia (hypergeometric test). **b.** Graphical representation from SYNGO of the synaptic compartment enrichment using membrane SEAD1 interactors list. **c.** SEAD1 in vivo injection site into mouse anterior cingulate cortex. Representative confocal image of SEAD1 and PSD95 immunofluorescence colocalization. **d.** Quantitation of 3D colocalization of SEAD1 with PSD95 vs. Gephyrin (N=3 mice, unpaired t-test, **: p<0.01). **e.** SEAD1 membrane interactors are mainly localized in VGLUT1+ synapses compared to VGAT+ synapses (One Way ANOVA, Tukey posthoc test).

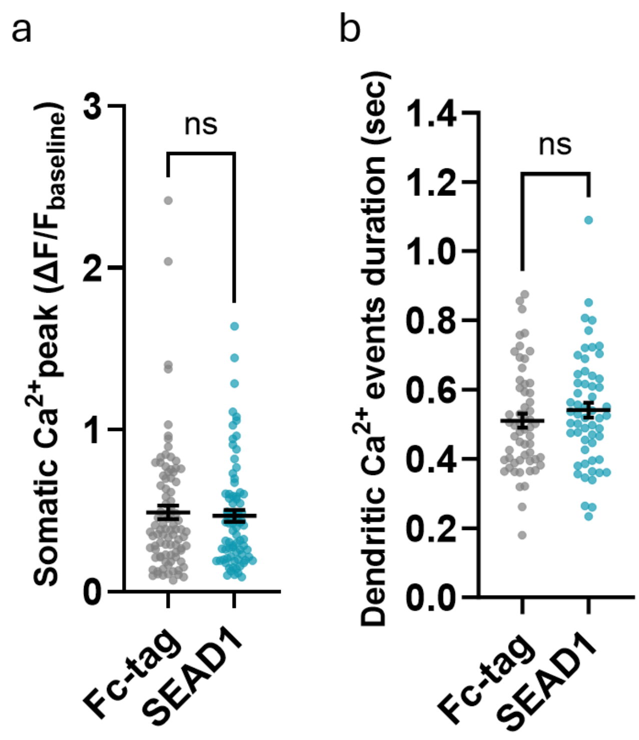


**Extended data Fig. 5. Supplementary metrics for effects of SEAD1 on dlx+ neurons $Ca^{2+}$ activity in vitro.**
**a.** Somatic Ca2+ events duration (t-test, **: p<0.01). (t-test, n.s., n=81/89 neurons). **b.** Dendritic Ca2+ event durations (t-test, n.s.).

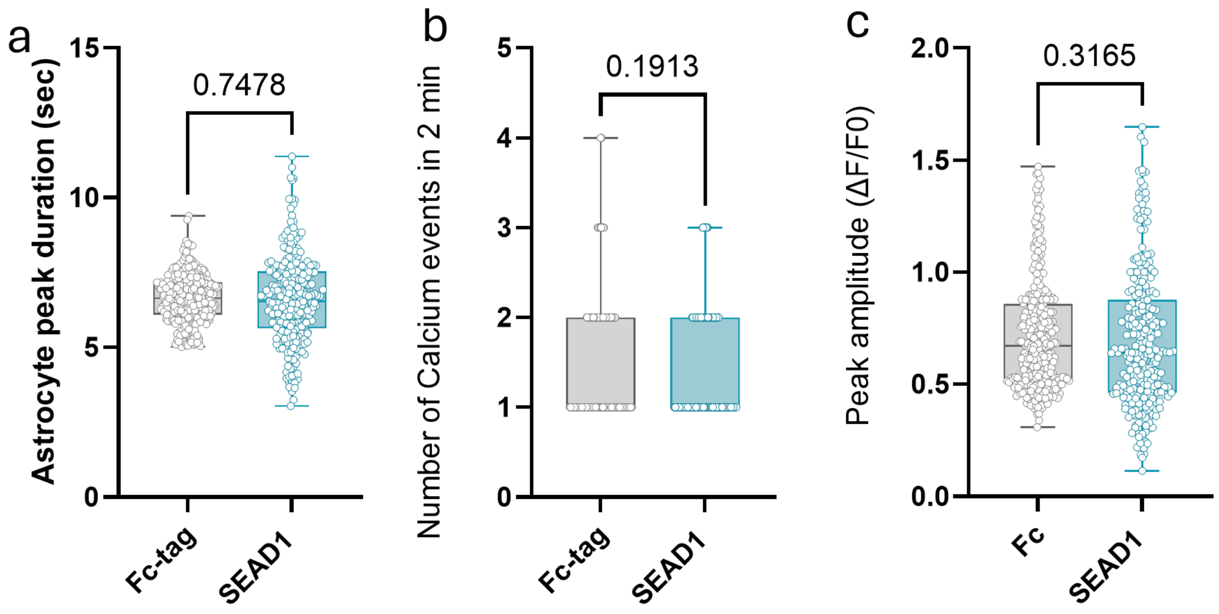


**Extended data Fig. 6. SEAD1 does not alter astrocytic calcium activity in vitro.** **a.** Measurement of the duration of astrocytic $Ca^{2+}$ in cortical neurons in culture reveals no difference between Fc-tag and SEAD1 conditions. **b.** Measurement of the number of astrocytic $Ca^{2+}$ in cortical neurons in culture reveals no difference between Fc-tag and SEAD1 conditions. **c.** Measurement of the amplitude of $Ca^{2+}$ in cortical neurons in culture reveals no difference between Fc-tag and SEAD1 conditions. T-test.

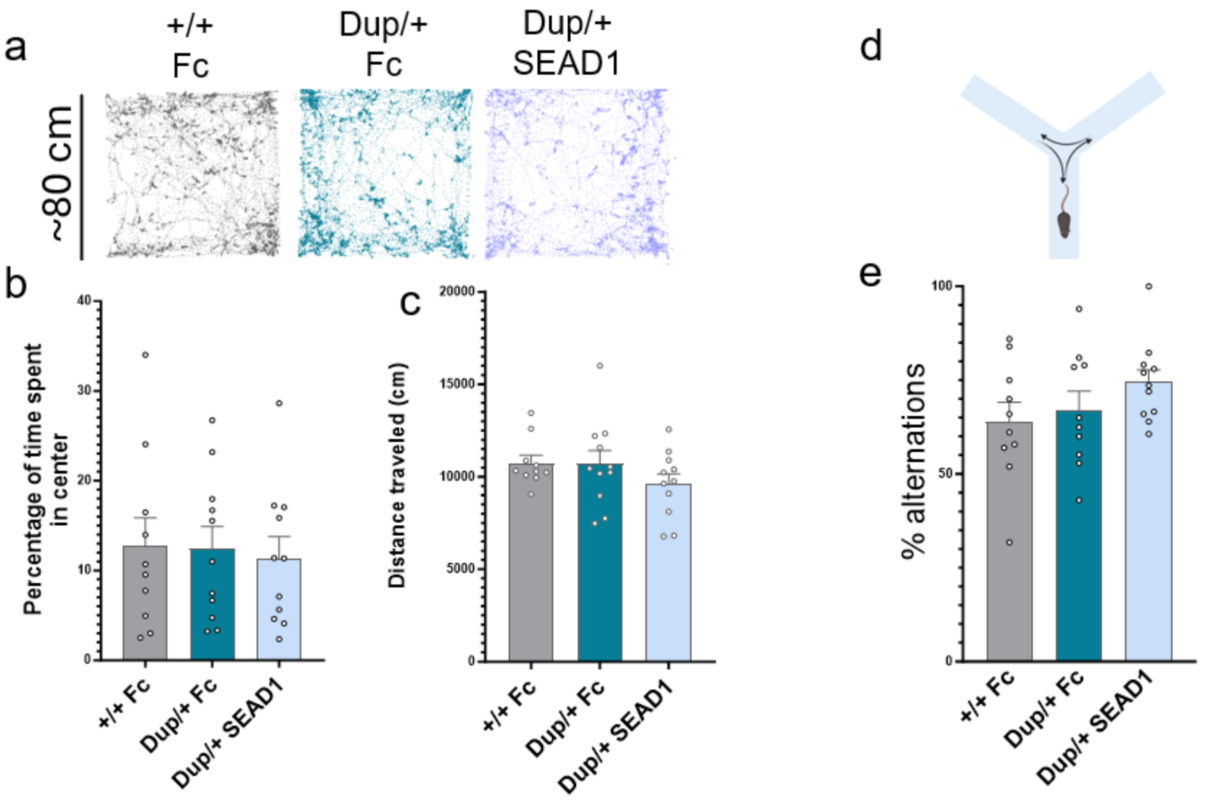


**Extended data Fig. 7. SEAD1 treatment does not alter basal locomotion, anxiety, or working memory.**

**a.** Representative centroids of the mice position for each frame of the 20-minute open field recording. **b.** The percentage of time spent in the center of the arena shows no difference between conditions. **c.** Distance traveled in 20 min is identical between conditions (Two-way ANOVA, no significance). d. Schematics of a Y-maze. **e.** The percentage of complete arm alternations is the same between conditions, no effect of genotype or treatment is observed.
